# Cross-sectional and longitudinal associations of family income-to-needs ratio with cortical and subcortical brain volume in adolescent boys and girls

**DOI:** 10.1101/2020.01.24.918847

**Authors:** Lucy S. King, Emily L. Dennis, Kathryn L. Humphreys, Paul M. Thompson, Ian H. Gotlib

**Author notes:** Corresponding author: Lucy S. King, Department of Psychology, Stanford University Jordan Hall, 450 Serra Mall, Building 420 Stanford, CA 94305, USA.

## Abstract

Deviations in neurodevelopment may underlie the association between lower childhood socioeconomic status and difficulties in cognitive and socioemotional domains. Most previous investigations of the association between childhood socioeconomic status and brain morphology have used cross-sectional designs with samples that span wide age ranges, occluding effects specific to adolescence. Sex differences in the association between childhood socioeconomic status and neurodevelopment may emerge or intensify during adolescence. We used tensor-based morphometry, a whole brain approach, to examine sex differences in the cross-sectional association between normative variation in family income-to-needs ratio (INR) and cortical and subcortical gray and white matter volume during early adolescence (ages 9-13 years, N=147), as well as in the longitudinal association between in INR and change in volume from early to later adolescence (ages 11-16 years, N=109). Biological sex interacted with INR to explain variation in volume in several areas cross-sectionally and longitudinally. Effects were primarily in cortical gray matter areas, including regions of the association cortex and sensorimotor processing areas. Effect sizes tended to be larger in boys than in girls. Biological sex may be an important variable to consider in analyses of the effects of family income on structural neurodevelopment during adolescence.

**Highlights:**

- Sex-specific associations of SES with neurodevelopment may emerge in adolescence.
- We used a whole-brain approach to examine gray and white matter volume.
- Sex interacted with SES to explain variation in volume across adolescence.
- Sex is an important variable to consider in analyses of SES and brain volume.

## 1. Introduction

Approximately 39% of children live in low-income households in the United States (defined by a family income below 200% of the poverty threshold; Semega, Fontenot and Kollar, 2017). Household income is one metric of socioeconomic status (SES), or an individual’s position in society in terms of material and non-material resources (e.g., education; Farah, 2017). From a measurement perspective, SES can be assessed on a continuum in which risk for negative outcomes is identified dimensionally in relation to relative resources, with the poorest children being at highest risk for experiencing difficulties in psychological functioning (Adler et al., 1994; Farah, 2018). While children in poverty may develop unique strengths that enhance adaptation to their environments (Frankenhuis & Nettle, 2019), exposure to lower SES in childhood is associated with, on average, poorer language ability (Fernald, Marchman, & Weisleder, 2013), executive function (Lawson, Hook, & Farah, 2018), and mental health (Amone-P’Olak, Burger, Huisman, Oldehinkel, & Ormel, 2011). Although it is not yet clear precisely *how* a relative lack of socioeconomic resources leads to deficits in these domains, researchers have theorized that SES affects neurodevelopment in a manner that leads to disparities in psychological functioning (Farah, 2017; Hackman, Farah, & Meaney, 2010).

A growing body of research indicates that SES is associated with variations in several aspects of brain structure. Previous studies have used whole-brain approaches to examine the associations between SES and cortical thickness and surface area. In large samples of children, adolescents, and young adults from the multi-site Pediatric Imaging, Neurocognition, and Genetics study, higher family income was associated with greater cortical surface area, with the strongest effects in brain regions that support language and executive function, including the bilateral inferior temporal, insula and inferior frontal gyri, and in the right occipital and medial prefrontal cortex (Brito & Noble, 2018; Brito, Piccolo, & Noble, 2017; Noble et al., 2015). In a another large sample of participants ages 5-25 years, there were positive associations between SES (family income and parental education) and cortical surface area in regions supporting sensorimotor processing, emotion regulation, language, and memory, including bilateral regions of the lateral prefrontal, anterior cingulate, lateral temporal, and superior parietal lobule (McDermott et al., 2019).

In addition to associations with cortical surface area, lower SES in childhood has been related to smaller cortical and subcortical brain volume. More specifically, lower family income has been associated with reduced total gray matter volume (Mackey et al., 2015; McDermott et al., 2019), and higher SES has been associated with larger hippocampal volume in samples spanning childhood to young adulthood (Ellwood-Lowe et al., 2018; Hanson, Chandra, Wolfe, & Pollak, 2011; Hanson et al., 2015; Jednoróg et al., 2012; McDermott et al., 2019; Noble, Houston, Kan, & Sowell, 2012; Yu et al., 2018) and, recently, with larger thalamus and striatum volume (McDermott et al., 2019). Finally, SES has been associated with white matter volume (McDermott et al., 2019; Ursache & Noble, 2016) and with the organization (i.e., the diffusion properties) of white matter tracts. Specifically, lower SES has been associated with lower fractional anisotropy in the superior longitudinal fasciculus (Rosen, Sheridan, Sambrook, Meltzoff, & McLaughlin, 2018), arcuate fasciculus (Noble, Korgaonkar, Grieve, & Brickman, 2013), and parahippocampal cingulum (Ursache & Noble, 2016). Overall, findings of extant research indicate that SES has widespread associations with brain structure.

The goal of the current study was to build on existing research to advance our understanding of the relation between SES and neurodevelopment in two important ways. The sample for the current study was recruited from San Francisco Bay Area communities where the cost of living, as well as income and education, are on average higher than they are in other geographic regions (https://data.census.gov/). We operationalized SES as a continuum of family income-to-needs ratio^1^ (INR), and our sample included more individuals at the high than at the low end of this continuum.

First, we focused on the association of INR with levels and changes in brain volume during the transition from early (ages 9-14) to later adolescence (ages 11-16), using tensor-based morphometry (TBM), a whole-brain approach, to simultaneously model both cortical and subcortical gray and white matter volume. Although investigations with large samples spanning wide age ranges have yielded important information concerning the average effects of SES on brain volume irrespective of developmental stage, a more focused analysis of adolescence is warranted. Neural plasticity during adolescence is second only to infancy; this heightened plasticity may render the brain especially vulnerable to variation in environmental input during adolescence (Gee & Casey, 2015). Conceptualizing adolescence as an “organizational” period for the brain (Gur & Gur, 2016), researchers have posited that neurodevelopment during adolescence underlies the acquisition of complex skills involving executive function that are important for wellbeing in adulthood (Aoki, Romeo, & Smith, 2017). Given the sensitivity of the adolescent brain and the potentially cascading consequences of deviations in neurodevelopment, it is important to characterize more precisely the associations between SES and brain volume during this period.

Second, the current study investigates potential sex-specific associations of SES with neurodevelopment during adolescence. Although studies of the neural correlates of SES typically “control” statistically for the biological sex of participants (i.e., by holding sex constant in predictive models), researchers rarely test sex as a moderator of SES and brain structure. The confluence of sex differences in neurodevelopment (Dennison et al., 2013; Gur & Gur, 2016; Wierenga, Langen, Oranje, & Durston, 2014; Wierenga, Sexton, Laake, Giedd, & Tamnes, 2018) and in sensitivity to environmental input (Humphreys et al., 2018; Jaffee, Caspi, Moffitt, Polo-Tomás, & Taylor, 2007; Whittle et al., 2014, 2017) may lead to sex differences in the associations of SES with brain structure. For example, the effects of environmental input are likely to depend on the nature of ongoing neurodevelopmental processes (Bock, Wainstock, Braun, & Segal, 2015), which are sexually dimorphic (i.e., whereas some regions develop faster in boys, others develop faster in girls; [Dennison et al., 2013; Gur & Gur, 2016; Wierenga et al., 2014]). Importantly, there is growing evidence that environmental input, including variation in SES, has sex-specific effects on brain structure. Whittle et al. (2014) found that adolescent boys were more sensitive than were girls to variation in environmental input in the form of positive caregiving, such that higher frequency of positive caregiving predicted attenuated volumetric growth in the amygdala and accelerated cortical thinning in the right anterior cingulate across adolescence in boys but not in girls. With respect to SES, Mcdermott et al. (2019) found that the positive association between SES and cortical surface area was significantly stronger in boys than in girls, whereas Kim et al. (2018) found that lower family income was associated with decreased structural brain network efficiency at ages 6-11 years in girls but not in boys. The current study builds on these findings by focusing on sex differences in the relation between SES and brain volume across the transition from early to later adolescence, a period in which sex differences may emerge or intensify (Gur & Gur, 2016).

## 2. Method

### 2.2 Participants

Participants were 214 adolescents and their parents who were recruited from Bay Area communities through local and media postings to participate in a longitudinal study of the psychobiological effects of early life stress (ELS) across the transition from early (Time 1[T1]) to later adolescence (Time 2[T2]; Humphreys, Kircanski, Colich, & Gotlib, 2016; King, Humphreys, Camacho, & Gotlib, 2019). Inclusion criteria were that the adolescents be between 9 and 13 years of age and be proficient in spoken English. In addition, adolescents were recruited such that the majority were in early puberty at T1 based on self-reported Tanner stage (96% self-reported Tanner stage ≤ 3; Marshall and Tanner, 1968; Morris and Udry, 1980); further, boys and girls were matched on the basis of self-reported Tanner staging rather than age. Self-reported Tanner stage is significantly positively associated with pubertal hormones in this sample (King, Graber, Colich, & Gotlib, 2020). Given sex differences in pubertal timing (Negriff & Susman, 2011), samples of adolescent boys and girls that are matched on age are confounded by sex differences in pubertal stage. Exclusion criteria at T1 included a history of major neurological or medical illnesses, severe learning disabilities that would affect comprehension of study procedures, presence of MRI contraindication (e.g., metal implants or braces), and, for girls, the onset of menses. Of the 214 adolescents who participated in the study at T1, 28 opted to not participate in the MRI session and 23 did not provide usable structural MRI (sMRI) data (e.g., due to movement during the scan). Among these participants, an additional 18 of their parents did not provide INR information, yielding a final T1 sample of 147 (57% female). At T2 (two years following T1), 111 adolescents participated in a second MRI session of whom 109 provided usable sMRI data. The participants with missing sMRI data did not differ significantly in INR or distribution of sex from those included here. Sample characteristics for adolescents included in the analyses are presented in Table 1.

**Table 1.**
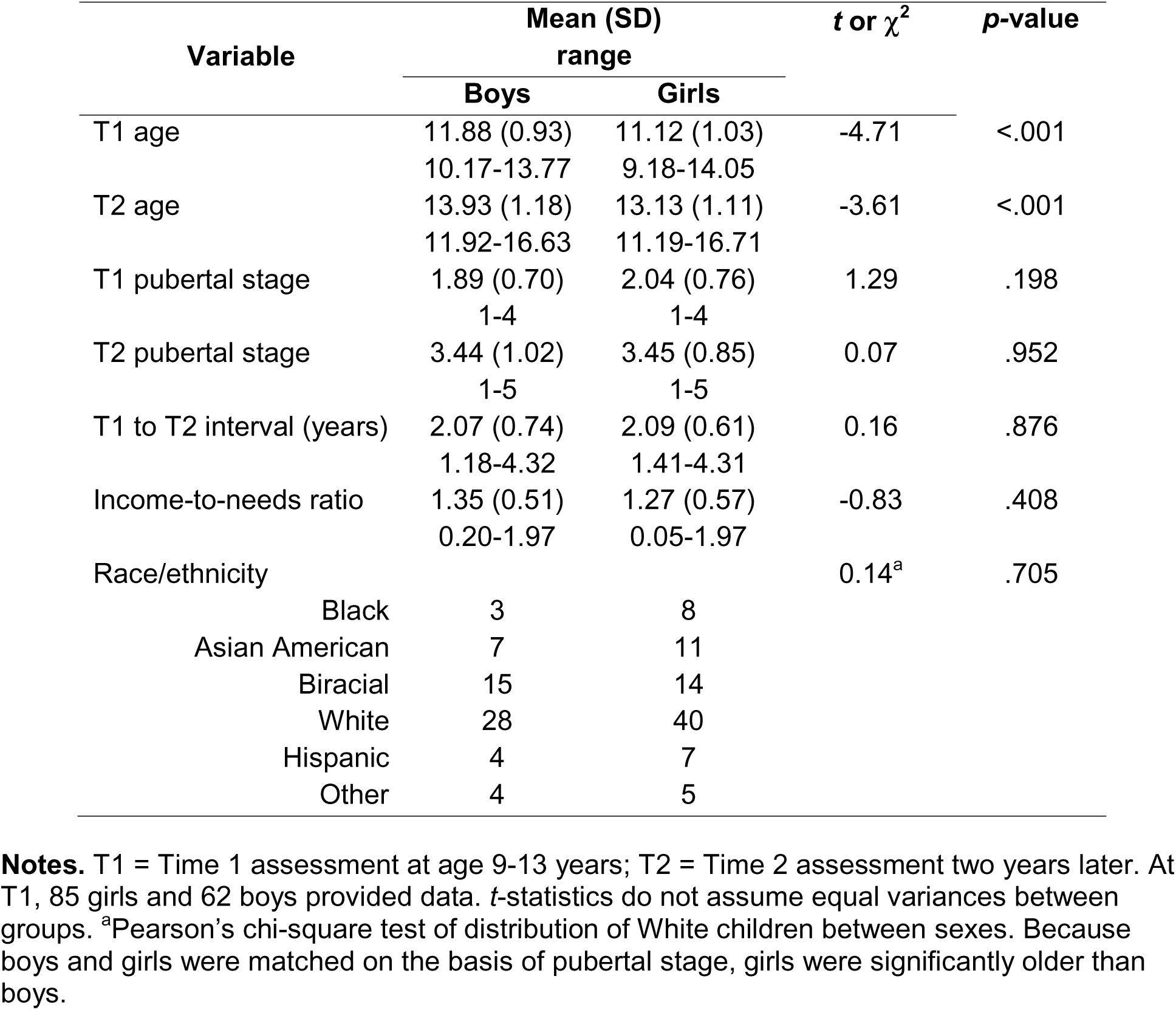
Sample characteristics

| Variable | Mean (SD)<br>range | | *t* or $\chi^2$ | *p*-value |
| --- | --- | --- | --- | --- |
|  | Boys | Girls |  |  |
| T1 age | 11.88 (0.93)<br>10.17-13.77 | 11.12 (1.03)<br>9.18-14.05 | -4.71 | <.001 |
| T2 age | 13.93 (1.18)<br>11.92-16.63 | 13.13 (1.11)<br>11.19-16.71 | -3.61 | <.001 |
| T1 pubertal stage | 1.89 (0.70)<br>1-4 | 2.04 (0.76)<br>1-4 | 1.29 | .198 |
| T2 pubertal stage | 3.44 (1.02)<br>1-5 | 3.45 (0.85)<br>1-5 | 0.07 | .952 |
| T1 to T2 interval (years) | 2.07 (0.74)<br>1.18-4.32 | 2.09 (0.61)<br>1.41-4.31 | 0.16 | .876 |
| Income-to-needs ratio | 1.35 (0.51)<br>0.20-1.97 | 1.27 (0.57)<br>0.05-1.97 | -0.83 | .408 |
| Race/ethnicity |  |  | 0.14<sup>a</sup> | .705 |
| Black | 3 | 8 |  |  |
| Asian American | 7 | 11 |  |  |
| Biracial | 15 | 14 |  |  |
| White | 28 | 40 |  |  |
| Hispanic | 4 | 7 |  |  |
| Other | 4 | 5 |  |  |

**Notes.** T1 = Time 1 assessment at age 9-13 years; T2 = Time 2 assessment two years later. At T1, 85 girls and 62 boys provided data. *t*-statistics do not assume equal variances between groups. <sup>a</sup>Pearson's chi-square test of distribution of White children between sexes. Because boys and girls were matched on the basis of pubertal stage, girls were significantly older than boys.

### 2.3 Procedure

The Stanford University Institutional Review Board approved the protocol for this study. In an initial telephone call, research staff provided information about the protocol to families and screened participants for inclusion/exclusion criteria. We then invited eligible families to attend a laboratory session during which staff obtained consent from parents and assent from adolescents. In this session, both parents and children completed interview and questionnaire measures about the adolescent and family. Adolescents completed the MRI scan at a follow-up session that occurred approximately 2 weeks following the initial laboratory visit. These procedures were repeated at a T2 session conducted an average of two years later (mean[SD]=2.08[0.67]; range: 1.18-4.32).

### 2.4 Measures

#### 2.4.1 Income-to-needs ratio

To operationalize SES, we calculated the INR for each child’s family. The parent who accompanied the child to the laboratory session reported total family income over the previous 12 months and the number of people in their family. We collected income data in bins, with parents reporting income on a 10-point scale as follows: <$5,000 (N=1); $5,001-$10,000 (N=2); $10,001-$15,000 (N=1); $15,001-$25,000 (N=6); $25,001-$35,000 (N=2); $35,001-$50,000 (N=10); $50,001-$75,000 (N=13); $75,000-$100,000 (N=17); $100,001-$150,000 (N=40); ≥$150,000 (N=54). To calculate the INR, we divided the midpoint of the endorsed income bin by the low-income limit for Santa Clara county (80% of the median income) determined by the Department of Housing and Urban Development based on the number of people in the household (https://www.huduser.gov/portal/datasets/il/il2017/2017summary.odn). Although collecting income data in bins, rather than asking for exact income, is a standard approach (McDermott et al., 2019; Noble et al., 2015), this approach may have truncated the distribution of INR in this study (i.e., families who reported incomes ≥$150,000 and had the same number of people in the household received equivalent values for INR despite likely variation above this threshold).

#### 2.4.2 MRI data acquisition

MRI scans were acquired at the Center for Cognitive and Neurobiological Imaging at Stanford University using a 3 T Discovery MR750 (GE Medical Systems, Milwaukee, WI, USA) equipped with a 32-channel head coil (Nova Medical, Wilmington, MA, USA). Whole-brain T1-weighted images (T1w) were collected using the following spoiled gradient echo pulse sequence: 186 sagittal slices; TR (repetition time)/TE (echo time)/TI (inversion time)=6.24/2.34/450ms; flip angle=12°; voxel sizeL=L0.9Lmm×0.9 mm×0.9 mm; scan duration=5:15 minutes.

#### 2.4.3 Tensor-based morphometry

Each participant’s T1-weighted anatomical data were N3-corrected using c3d (http://www.itksnap.org) to correct for intensity inhomogeneities. Volumes were automatically skull-stripped using Brainsuite and brain masks were manually edited to remove extraneous skull or meninges by trained MRI research coordinators (LS, AC, and AO, see Acknowledgements). We linearly registered each participant to the Montréal Neurological Institute (MNI) template using FSL FLIRT (http://fsl.fmrib.ox.ac.uk) with concatenated 6, 7, 9, and 12 degree of freedom (DOF) transformations. Thirty participants, selected to be representative of the population, were used to make the minimal deformation template (MDT). The MDT is the template that deviates least from the anatomy of the participants with respect to a mathematically defined metric of difference; in some circumstances, using an MDT can improve statistical power (Leporé et al., 2007). The MDT serves as an unbiased registration target for nonlinear registrations.

Next, each participant’s masked, non-uniformity-corrected, template-aligned T1-weighted image was non-linearly aligned to the MDT, using Advanced Normalization Tools Symmetric Normalization (SyN; Avants et al., 2008). SyN registration utilized a multi-level approach, i.e., the “moving” and fixed T1 images were successively less smoothed at each level, with a full resolution registration occurring at the final level. We used 150, 80, 50, and 10 iterations at each level, with a Gaussian kernel smoothing sigma set to 3, 2, 1, and 0 respectively (7.05, 4.7, 2.35, and 0 voxels full width at half maximum). Image similarity was measured using the ANTs implementation of mutual information (Avants et al., 2011). Image intensities were winsorized, excluding top and bottom 1% of voxels, and histogram matching was used. This process resulted in jacobian determinant images, where the values indicate the direction and magnitude of the deformation in registering an individual’s T1 to the MDT. For longitudinal analyses, each participant’s template-aligned T1 from the follow-up scan was non-linearly aligned to the template-aligned T1 from the first scan using the same parameters listed above. The output jacobian determinant image showed the direction and magnitude of the change between the participant’s T1 and T2 anatomical images. Data were quality checked at multiple stages, including brain masking, after linear registration, and after non-linear registration.

### 2.5 Statistical Analyses

In all analyses, we modeled INR as a square-root term to capture the potential asymptotic relation between income and brain structure previously reported (Hair, Hanson, Wolfe, & Pollak, 2015; Noble et al., 2015), such that the steepest gradient is at the lower end of the INR continuum. Analysis scripts are available at https://github.com/lucysking/income_TBM. Raw data from the study are uploaded to the NIMH RDoC repository twice per year (https://data-archive.nimh.nih.gov/). Data specific to the current analyses are available upon request.

#### 2.5.1 Sex differences in the association between INR and brain volume

To examine sex differences in the cross-sectional association between INR and gray and white matter volume in early adolescence and longitudinal changes in gray and white matter volume from early to later adolescences, we conducted separate voxel-wise linear regressions testing the *interactive* effects of sex and INR on brain volume. The 9-DOF linear registration that is part of the TBM processing protocol accounts for differences in overall brain scale, removing much of the effect of intracranial volume (ICV); nevertheless, we also included ICV, computed from the linearly registered image, as a covariate. We present the cross-sectional and longitudinal models in equations 1 and 2, respectively:

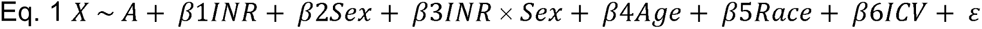

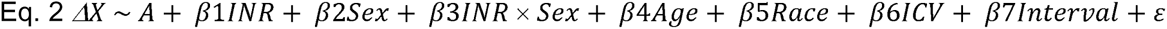

where *X* is the jacobian determinant value at a given position in early adolescence, Δ*X* is the change in volume from early to later adolescence, *A* is the constant jacobian determinant term, the βs are the regression coefficients, and *ϵ* is an error term. For each model, results were corrected for multiple comparisons across all voxels tested using Searchlight FDR (Langers, Jansen, & Backes, 2007). We report only those clusters that exceeded 50 voxels. Because each model examined a distinct dependent variable (cross-sectional volume versus longitudinal change in volume), we treated each as a separate hypothesis test and did not adjust for multiple tests at the model level.

#### 2.5.2 Simple effects of INR on brain volume within each sex

To characterize significant INR × sex interactions (i.e., to determine the strength and the direction of the association of INR with brain volume within each sex), we extracted the estimated volumes of each significant cluster identified in the cross-sectional and longitudinal whole-brain analyses. Using the statistical program R (R Core Team, 2018), we then conducted separate linear regression models for each extracted cluster as follows in equation 3:

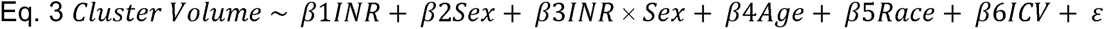

From the results of each model, we computed the standardized betas and 95% confidence intervals for the simple effects of INR in boys and girls, respectively.

#### 2.5.3 Functional correlates of anatomical results

Following previous research using a whole-brain approach to examine effects of variation in childhood SES (McDermott et al., 2019), the coordinates of the cluster peaks in gray matter identified in the primary analyses were submitted to Neurosynth (Yarkoni, Poldrack, Nichols, Essen, & Wager, 2011), which provides empirically established mappings between psychological or cognitive states and localized blood-oxygenation-level dependent (BOLD) activation. For each cluster, we identified the top three psychological or cognitive terms with the highest meta-analytic coactivation correlation coefficient after anatomical and redundant terms were removed. This method aided in interpreting the potential functional implications of the anatomical results.

## 3. Results

### 3.1 Sample characteristics

Sample characteristics are presented in Table 1. Based on having an INR<1.00, 29% of the sample was “low income.” Therefore, INR was negatively skewed, and results of our analyses are not necessarily generalizable to samples that do not include higher SES adolescents or to samples including a larger number of adolescents below the federal poverty line (annual income of ∼$25,000 for a family of four within the study period, equivalent to an INR of 0.29 in this study). Instead, our results may be interpreted as associations of INR with brain volume in the context of variation in relative socioeconomic enrichment. Boys and girls did not differ in pubertal stage at T1 or T2, the interval between T1 and T2, percentage White, INR, or threat-related ELS; however, because boys and girls were matched on pubertal stage at T1, girls were significantly younger than boys.

### 3.2 Sex differences in the association between INR and brain volume

There were several clusters with a significant interaction between INR and sex on brain volume cross-sectionally. Specifically, sex and INR interacted to explain brain volume in the midline cerebellar vermis, midbrain, bilateral hippocampal cingulum, thalamus, left lateral occipital gyrus, left inferior frontal gyrus, left angular gyrus, left fusiform gyrus, right posterior thalamic radiation, right postcentral gyrus, right superior and middle frontal gyrus, and right superior temporal gyrus. Longitudinally, sex and INR interacted to explain brain volume in the bilateral lingual gyrus, bilateral superior frontal gyrus, left inferior temporal gyrus, left cerebellar grey matter, left superior longitudinal fasciculus, right superior parietal lobule, right posterior thalamic radiation, and right hippocampal cingulum. The statistics for these results as well as the top three psychological terms from Neurosynth (Yarkoni et al., 2011) with the highest meta-analytic coactivation correlation coefficient are presented in **Table 2** and visualizations of the significant clusters are presented in **Figure 1**.

**Figure 1.**
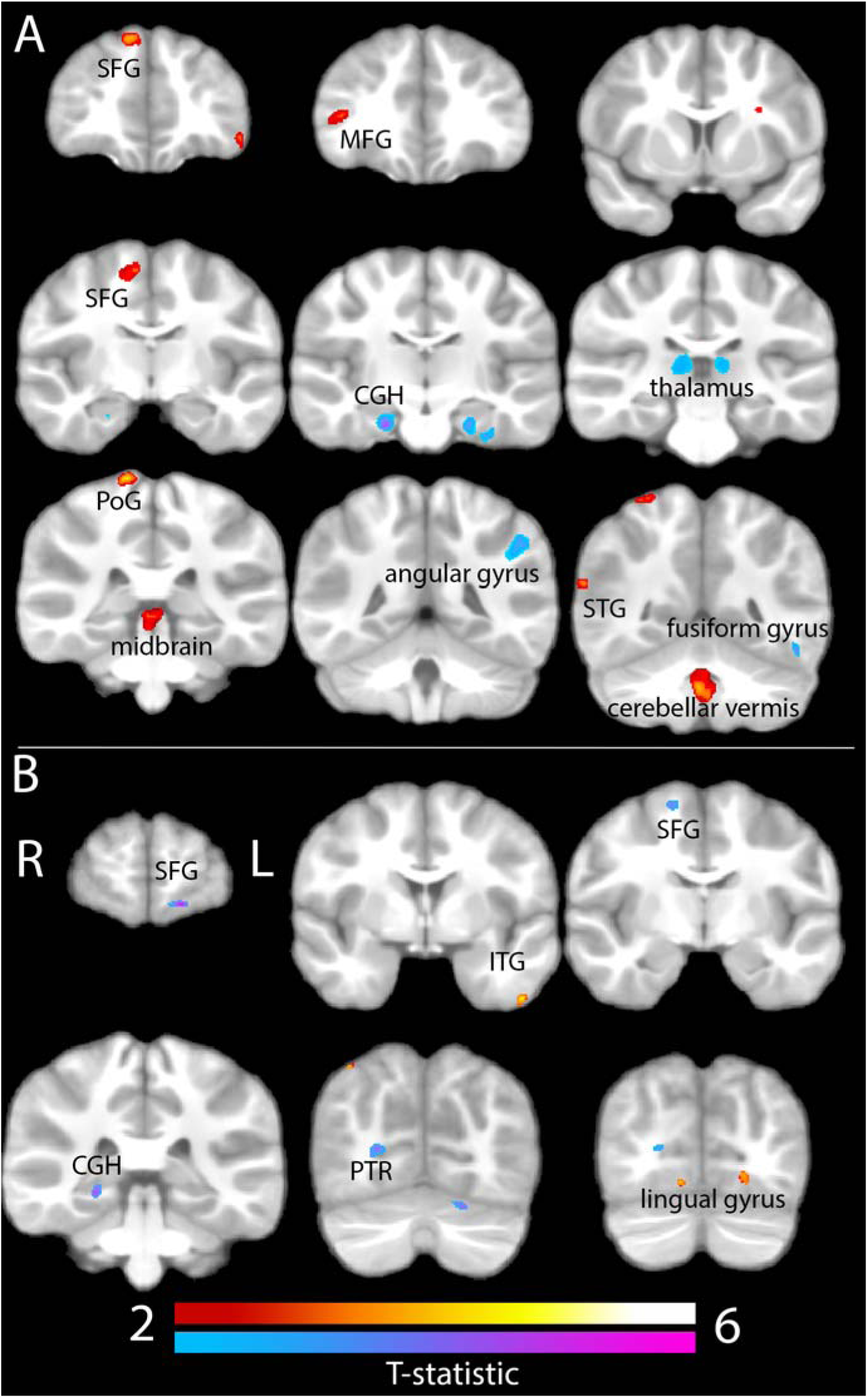
Cross-sectional (A) and longitudinal (B) results from the INR × sex analysis. **Notes**. Structure labeled based on location of cluster peak. Cross-sectional clusters (A): SFG = superior frontal gyrus, MFG = middle frontal gyrus, CGH = hippocampal cingulum, PoG = postcentral gyrus, STG = superior temporal gyrus. Longitudinal clusters (B): ITG = inferior temporal gyrus, PTR = posterior thalamic radiation. “Positive” (red scale) interactions indicate that the association between INR and volume is significantly more positive in boys than in girls; “negative” (blue scale) interactions indicate the association between INR and volume is significantly more negative in boys than in girls.

**Table 2.** Cross-sectional (A) and longitudinal (B) results from the INR × sex analysis.

| Voxels | MAX T-stat | X | Y | Z | Side | Structure | Tissue | Top Neurosynth terms |
| --- | --- | --- | --- | --- | --- | --- | --- | --- |
| **A. Cross-sectional** |  |  |  |  |  |  |  |  |
| **Positive** |  |  |  |  |  |  |  |  |
| 1831 | 4.17 | 3 | -47 | -35 | R | Cerebellar vermis | GM | encoding, memories, autobiographical |
| 739 | 3.41 | 3 | -32 | -4 | R | Midbrain | WM | N/A |
| 314 | 3.42 | 40 | -55 | 2 | R | Posterior thalamic radiation | WM | N/A |
| 288 | 3.54 | 9 | -8 | 54 | R | Superior frontal gyrus | WM | N/A |
| 268 | 4.39 | 15 | -32 | 69 | R | Postcentral gyrus | GM | motor, sensorimotor, somatosensory |
| 252 | 4.05 | 11 | 46 | 45 | R | Superior frontal gyrus | GM | mental states, mentalizing, theory of mind |
| 248 | 3.52 | 51 | 38 | 5 | R | Middle frontal gyrus | GM | retrieval, emotional, semantic |
| 162 | 3.64 | -23 | -89 | 3 | L | Lateral occipital gyrus | GM | visual, objects, reading |
| 130 | 3.6 | -48 | 46 | -8 | L | Inferior frontal gyrus | GM | semantic, retrieval, word |
| 106 | 3.86 | 63 | -47 | 19 | R | Superior temporal gyrus | GM | theory of mind, belief, intentions |
| **Negative** |  |  |  |  |  |  |  |  |
| 1056 | 3.9 | -46 | -39 | 33 | L | Angular gyrus | WM | N/A |
| 576 | 3.34 | 13 | -23 | 8 | R | Thalamus | GM | pain, motor, movement |
| 529 | 3.7 | -29 | -19 | -33 | L | Hippocampal cingulum/ Fusiform gyrus | WM | N/A |
| 513 | 4.91 | 23 | -14 | -23 | R | Hippocampal cingulum | WM | N/A |
| 373 | 3.19 | -9 | -24 | 10 | L | Thalamus | GM | goal, motor, pain |
| 192 | 3.93 | -49 | -49 | -16 | L | Fusiform gyrus | GM | visual word, word form, orthographic |
| **A. Longitudinal** |  |  |  |  |  |  |  |  |
| **Positive** |  |  |  |  |  |  |  |  |
| 80 | 4.04 | -20 | -75 | -5 | L | Lingual gyrus | WM | N/A |
| 79 | 4.54 | -48 | 1 | -40 | L | Inferior temporal gyrus | GM | theory of mind, social, mentalizing |
| 67 | 4.31 | 12 | -73 | -9 | R | Lingual gyrus | GM | visual, eye movements, sighted |
| 57 | 4.42 | 27 | -60 | 56 | R | Superior parietal lobule | GM | visual, spatial, tasks |
| **Negative** |  |  |  |  |  |  |  |  |
| 430 | 4.56 | 24 | -70 | 7 | R | Posterior thalamic radiation | WM | N/A |
| 346 | 4.58 | -18 | -66 | -21 | L | Cerebellum | GM | motor, production, movement |
| 168 | 3.96 | 17 | -5 | 58 | R | Superior frontal gyrus | WM | N/A |
| 160 | 5.28 | -16 | 62 | -10 | L | Superior frontal gyrus | GM | default mode, autobiographical, episodic |
| 96 | 4.16 | -33 | -44 | 16 | L | Superior longitudinal fasciculus | WM | N/A |
| 84 | 4.54 | 26 | -33 | -7 | R | Hippocampal cingulum | WM | N/A |

**Notes.** WM (white matter) vs. GM (gray matter) labeled based on location of cluster peak. Sex was dummy coded (1 = male; 0 = female) in the whole-brain analyses. “Positive” interactions indicate that the association between INR and volume is significantly more positive in boys than in girls; “negative” interactions indicate the association between INR and volume is significantly more negative in boys than in girls.

### 3.3 Simple effects of INR on brain volume within each sex

Results of linear regression analyses probing each significant INR × sex interaction identified in the cross-sectional and longitudinal whole-brain analyses are presented in Table 3. In both cross-sectional and longitudinal analyses, effect sizes tended to be larger in boys than in girls. This was particularly apparent in the longitudinal analyses, in which βs in boys ranged from +/- 0.51-0.90, whereas βs in girls ranged from +/- 0.02-0.19. Indeed, whereas INR was associated with volume in both boys and girls cross-sectionally, results suggested that INR was associated with *change* in volume from early to later adolescence in boys only. Cross-sectionally, most of the interactions were characterized by opposing (as opposed to weaker or stronger) effects in boys compared to girls. Generally, when INR was positively associated with volume in boys, it was at least weakly negatively associated with volume in girls. Several of the cross-sectional interactions were “reversal” interactions, such that effect sizes were nearly equivalent but in opposite directions in boys and girls. We visualize the simple effects in boys and girls in Figures 2 and 3.

**Figure 2.**
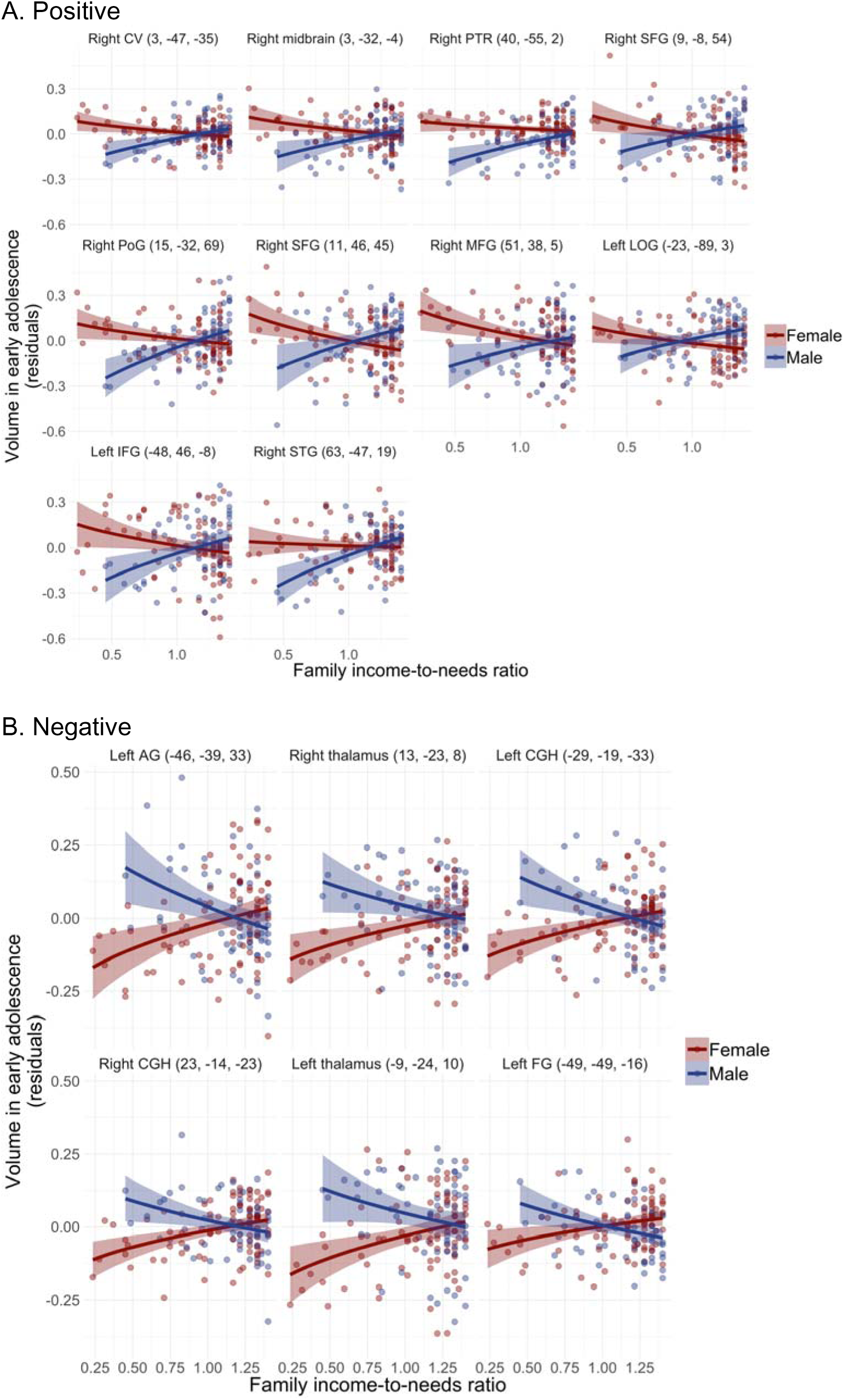
Scatterplots of results from cross-sectional INR × sex analysis in early adolescence. **Notes.** Coordinates of cluster peaks are in parentheses. Positive clusters (A): CV = cerebellar vermis, WM = white matter, PTR = posterior thalamic radiation, SFG = superior frontal gyrus, PoG = postcentral gyrus, MFG = middle frontal gyrus, LOG = lateral occipital gyrus, IFG = inferior frontal gyrus, STG = superior temporal gyrus. Negative clusters (B): AG = angular gyrus, CGH= hippocampal cingulum, FG = fusiform gyrus.

**Figure 3.**
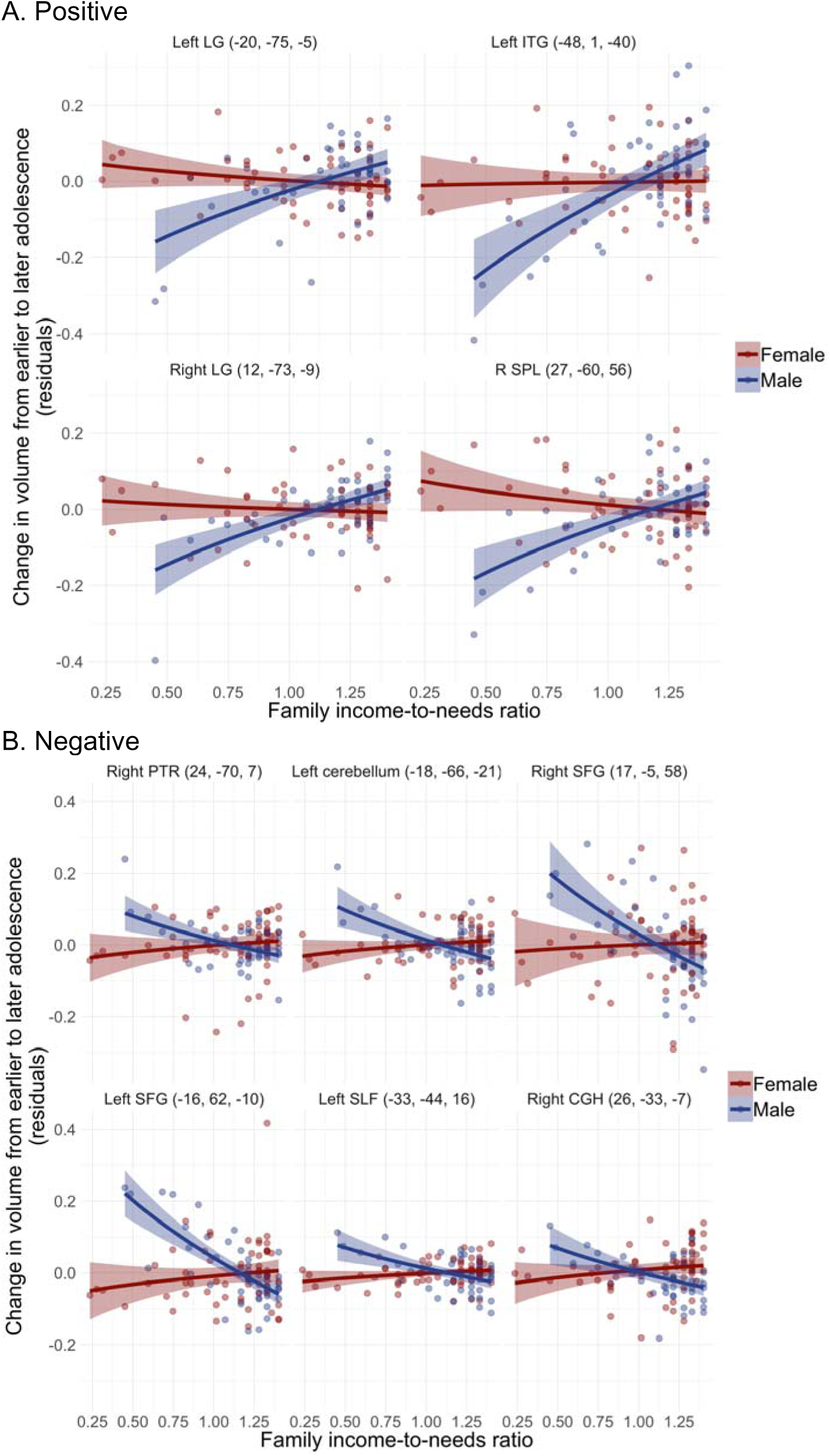
Scatter plots of results from longitudinal INR × sex analysis. **Notes.** Coordinates of clusters peaks are in parentheses. Positive clusters: LG = lingual gyrus, ITG = inferior temporal gyrus, SPL = superior parietal lobule. Negative clusters: PTR = posterior thalamic radiation, SFG = superior frontal gyrus, SLF = superior longitudinal fasciculus, CGH = hippocampal cingulum

**Table 3.**
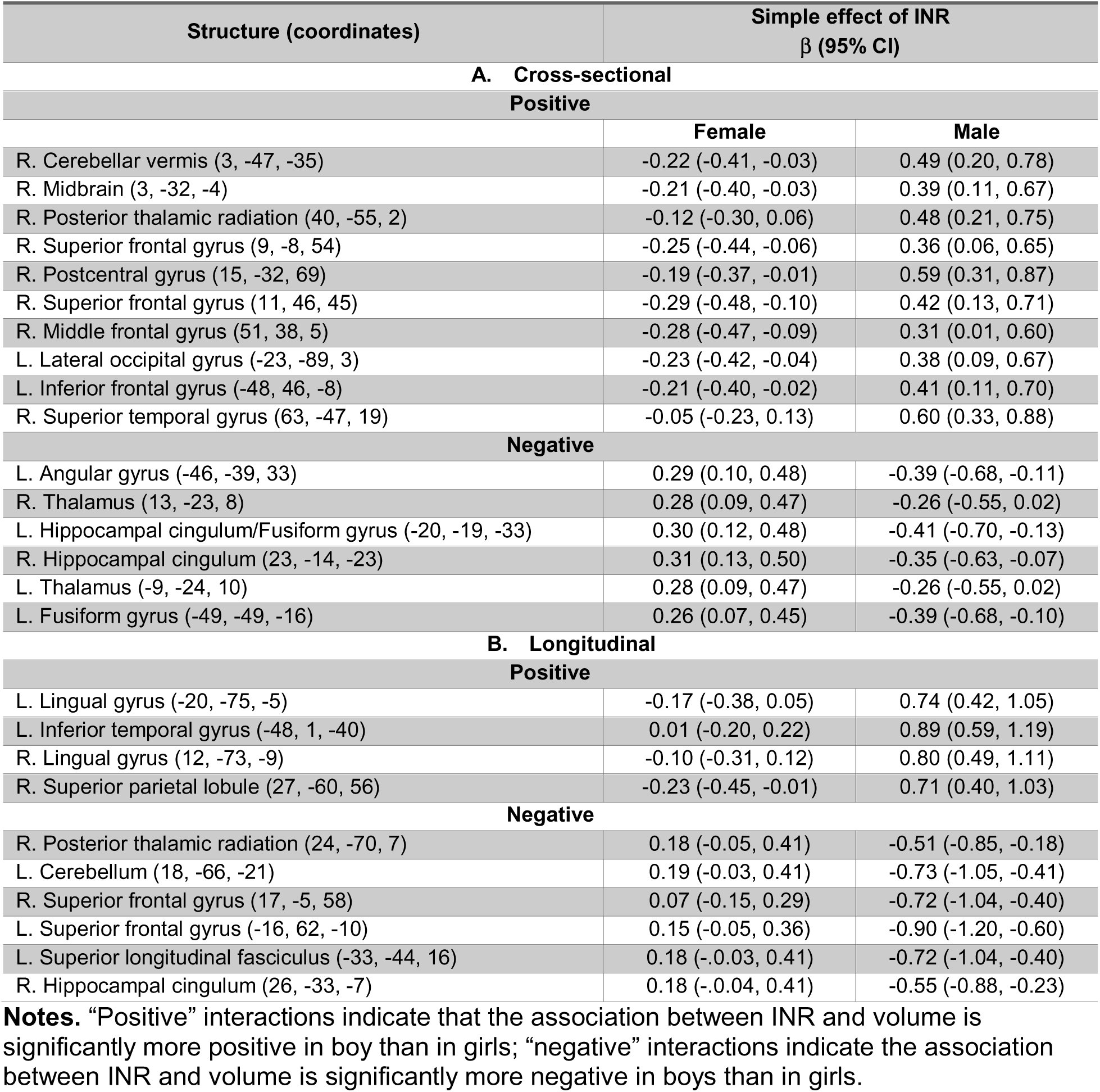
Simple effect sizes of cross-sectional (A) and longitudinal (B) associations of INR in boys and girls

| Structure (coordinates) | Simple effect of INR<br>β (95% CI) |  |
| --- | --- | --- |
| A. Cross-sectional |  |  |
| Positive |  |  |
|  | Female | Male |
| R. Cerebellar vermis (3, -47, -35) | -0.22 (-0.41, -0.03) | 0.49 (0.20, 0.78) |
| R. Midbrain (3, -32, -4) | -0.21 (-0.40, -0.03) | 0.39 (0.11, 0.67) |
| R. Posterior thalamic radiation (40, -55, 2) | -0.12 (-0.30, 0.06) | 0.48 (0.21, 0.75) |
| R. Superior frontal gyrus (9, -8, 54) | -0.25 (-0.44, -0.06) | 0.36 (0.06, 0.65) |
| R. Postcentral gyrus (15, -32, 69) | -0.19 (-0.37, -0.01) | 0.59 (0.31, 0.87) |
| R. Superior frontal gyrus (11, 46, 45) | -0.29 (-0.48, -0.10) | 0.42 (0.13, 0.71) |
| R. Middle frontal gyrus (51, 38, 5) | -0.28 (-0.47, -0.09) | 0.31 (0.01, 0.60) |
| L. Lateral occipital gyrus (-23, -89, 3) | -0.23 (-0.42, -0.04) | 0.38 (0.09, 0.67) |
| L. Inferior frontal gyrus (-48, 46, -8) | -0.21 (-0.40, -0.02) | 0.41 (0.11, 0.70) |
| R. Superior temporal gyrus (63, -47, 19) | -0.05 (-0.23, 0.13) | 0.60 (0.33, 0.88) |
| Negative |  |  |
| L. Angular gyrus (-46, -39, 33) | 0.29 (0.10, 0.48) | -0.39 (-0.68, -0.11) |
| R. Thalamus (13, -23, 8) | 0.28 (0.09, 0.47) | -0.26 (-0.55, 0.02) |
| L. Hippocampal cingulum/Fusiform gyrus (-20, -19, -33) | 0.30 (0.12, 0.48) | -0.41 (-0.70, -0.13) |
| R. Hippocampal cingulum (23, -14, -23) | 0.31 (0.13, 0.50) | -0.35 (-0.63, -0.07) |
| L. Thalamus (-9, -24, 10) | 0.28 (0.09, 0.47) | -0.26 (-0.55, 0.02) |
| L. Fusiform gyrus (-49, -49, -16) | 0.26 (0.07, 0.45) | -0.39 (-0.68, -0.10) |
| B. Longitudinal |  |  |
| Positive |  |  |
| L. Lingual gyrus (-20, -75, -5) | -0.17 (-0.38, 0.05) | 0.74 (0.42, 1.05) |
| L. Inferior temporal gyrus (-48, 1, -40) | 0.01 (-0.20, 0.22) | 0.89 (0.59, 1.19) |
| R. Lingual gyrus (12, -73, -9) | -0.10 (-0.31, 0.12) | 0.80 (0.49, 1.11) |
| R. Superior parietal lobule (27, -60, 56) | -0.23 (-0.45, -0.01) | 0.71 (0.40, 1.03) |
| Negative |  |  |
| R. Posterior thalamic radiation (24, -70, 7) | 0.18 (-0.05, 0.41) | -0.51 (-0.85, -0.18) |
| L. Cerebellum (18, -66, -21) | 0.19 (-0.03, 0.41) | -0.73 (-1.05, -0.41) |
| R. Superior frontal gyrus (17, -5, 58) | 0.07 (-0.15, 0.29) | -0.72 (-1.04, -0.40) |
| L. Superior frontal gyrus (-16, 62, -10) | 0.15 (-0.05, 0.36) | -0.90 (-1.20, -0.60) |
| L. Superior longitudinal fasciculus (-33, -44, 16) | 0.18 (-0.03, 0.41) | -0.72 (-1.04, -0.40) |
| R. Hippocampal cingulum (26, -33, -7) | 0.18 (-0.04, 0.41) | -0.55 (-0.88, -0.23) |

**Notes.** “Positive” interactions indicate that the association between INR and volume is significantly more positive in boy than in girls; “negative” interactions indicate the association between INR and volume is significantly more negative in boys than in girls.

#### 3.3.1 Sensitivity analyses

First, we conducted diagnostic tests to identify influential cases. We identified outlying observations in three of the regression models. When these outlying observations were removed, effect sizes were highly similar (detailed information is provided in the Supplementary Material). Second, given that girls were significantly younger than boys, we conducted sensitivity analyses in an age-matched subsample of 53 adolescents (24 girls) who were ages 11.00-12.00 years at T1. We reran the regression models probing each significant INR × sex interaction in this subsample, examining whether the strength and direction of effects within each sex were similar to those in the full sample. We found remarkably similar results in the subsample, which we present in the Supplementary Material (see Table S1 and Figures S1 and S2). For the majority of clusters, effect sizes (βs) in the age-matched subsample were similar to and in the same direction as those in the full sample, indicating that INR × sex interactions were not driven by sex differences in age. Cross-sectional associations between INR and volume in the left angular gyrus were attenuated in both boys and girls in the age-matched subsample. Further, whereas INR was not associated with change in volume in the right lingual gyrus in girls in the full sample, associations between INR and change volume in right lingual gyrus were moderately positive in both boys and girls in the subsample.

## 4. Discussion

The goal of the current study was to build on existing research to advance our understanding of the relation between SES and neurodevelopment. Specifically, using tensor-based morphometry (TBM) we extended previous research by investigating sex-specific associations of family income-to-needs ratio (INR) with cortical and subcortical gray and white volume in early adolescence (ages 9-13 years), and with changes in volume from early to later adolescence (ages 11-16 years). We found that sex interacted with INR to explain brain volume in a number of regions, including in areas of the association cortex, sensorimotor processing areas, the thalamus, the hippocampal cingulum, and the superior longitudinal fasciculus. Overall, our findings replicate those of previous studies that have identified widespread associations of SES with brain structure (McDermott et al., 2019; Noble et al., 2015) but suggest that during adolescence the nature of these associations depend on biological sex. Adolescents in the current study were recruited from San Francisco Bay Area communities where the cost of living as well as income and education are higher than they are in other geographic regions (https://data.census.gov/). Although 29% of adolescents were designated “low-income” based on their family INR, there were more adolescents at the higher end of the INR continuum than at the lower end. Therefore, results of the current study may be best interpreted as the associations of normative variation in INR with brain volume. Broadly, variation in INR was more strongly associated with brain volume in boys than in girls.

Consistent with several previous investigations of the associations of socioeconomic status (SES) with gray matter morphology in samples spanning wide age ranges (Brito & Noble, 2018; Brito et al., 2017; McDermott et al., 2019; Noble et al., 2015), we found that INR was associated with volume in several areas of the association cortex. Specifically, we found that cortical regions associated with language, mentalizing, theory of mind, and emotion (Yarkoni et al., 2011)—including areas of the fusiform gyrus, inferior frontal gyrus, middle frontal gyrus, superior frontal gyrus, and inferior and superior temporal gyri—were sensitive to INR in interaction with biological sex in early adolescence and/or longitudinally from early to later adolescence. Simple effects analyses indicated that the association between INR and volume in areas of the association cortex tended to be stronger in boys than in girls both cross-sectionally and longitudinally. In early adolescence, INR was moderately positively associated with volume in these areas in boys whereas INR was weakly negatively associated with volume in these areas in girls. Across the transition from early to later adolescence, INR was not associated with changes in volume in areas of the association cortex in girls. However, in boys INR was strongly associated with *greater* volume expansion in an area of the left inferior temporal gyrus implicated in theory of mind, and was strongly associated with *less* volume expansion, or more contraction, in an area of the bilateral superior frontal gyrus implicated in the default mode.

In addition to effects in areas of the association cortex, we found that INR interacted with biological sex both cross-sectionally and longitudinally to explain variation in volume in regions of the cortex involved in motor function and sensory processing, including in areas of the postcentral gyrus, lateral occipital gyrus, fusiform gyrus, superior parietal lobule, and lingual gyrus (Yarkoni et al., 2011). It is noteworthy that in a recent study using an accelerated longitudinal design spanning early childhood to young adulthood, the effects of childhood SES on cortical surface area and thickness were also localized to regions associated with sensorimotor processing (McDermott et al., 2019). In the current study, simple effect analyses indicated that effect sizes within these regions were similar in boys and girls in early adolescence, but in opposite directions. Specifically, higher INR was associated with larger volume in the right postcentral gyrus and left lateral occipital gyrus in boys and smaller volume in these areas in girls. In contrast, in the left fusiform gyrus, higher INR was associated with larger volume in girls and smaller volume in boys. Across the transition from early to later adolescence, higher INR was strongly associated with greater volume expansion in the bilateral lingual gyrus and in the right superior parietal lobule in boys; in girls, however, INR was weakly associated with less volume expansion, or more contraction, in these regions.

Although most of the associations that we found were in cortical regions, we also found that INR interacted with sex to explain variation in volume of the bilateral thalamus in early adolescence. This is the second recent study to identify an association of childhood SES with thalamic volume (McDermott et al., 2019), a sensory relay region. Activation of the areas of the thalamus we identified is associated meta-analytically with the processing of pain, movement, and goal-related behavior (Yarkoni et al., 2011). In both boys and girls, effect sizes for the association between INR and thalamic volume were in the small to medium range, but in opposite directions. In boys, INR was negatively associated with bilateral thalamic volume whereas in girls INR was positively associated with bilateral thalamic volume.

Given the central role of the hippocampus in memory, learning, and regulation of responses to environmental stress (Teicher & Samson, 2016), several previous studies of SES and neurodevelopment have examined hippocampal volume using a region-of-interest approach (Ellwood-Lowe et al., 2018; Hanson et al., 2011, 2015; Jednoróg et al., 2012; McDermott et al., 2019; Noble et al., 2012; Yu et al., 2018). In addition, researchers have examined the relation between SES and microstructure of the hippocampal subdivision of the cingulum, which, extending into the temporal lobe, is implicated in memory performance (Bubb, Metzler-Baddeley, & Aggleton, 2018). Specifically, in a large sample spanning childhood to adulthood, Ursache and Noble (2016) found that higher family income was associated with greater fractional anisotropy—a measure of the degree of fiber coherence or directionality—in the right hippocampal cingulum. In the current study, INR interacted with sex to explain variation in volume of the bilateral hippocampal cingulum^2^ in early adolescence and in change in volume of the right hippocampal cingulum from early to later adolescence. In early adolescence, higher INR was moderately associated with *smaller* volume in the bilateral hippocampal cingulum in boys and moderately associated with *larger* volume in these areas in girls. Longitudinally, associations were again negative in boys and positive in girls: higher INR was moderately associated with less volume expansion, or greater contraction, in the right hippocampal cingulum in boys, but was weakly associated with greater volume expansion in girls.

Our findings increase specificity in our knowledge of the effects of SES on neurodevelopment by characterizing the association of INR with brain volume during adolescence, a period of high neural plasticity in which environmental input may have outsized effects on development. In addition, in contrast to most previous research in which sex has been treated as a covariate rather than as a moderator, we examined sex-specific associations of INR with brain volume. Our finding that cross-sectional and longitudinal associations of INR with brain volume depended on biological sex may be explained by aspects of our study design. Sex differences in brain organization that are evident in adults intensify in adolescence, likely because of hormonal changes associated with puberty (Gur & Gur, 2016). The broader study from which the current data are drawn was specifically designed to examine sex differences in brain development in adolescence; in fact, given theory and evidence that neuroplasticity during adolescence is driven by pubertal maturation (Herting et al., 2014; Wierenga, Bos, et al., 2018), we matched boys and girls at entry to the study based on pubertal stage. Therefore, unlike studies in which boys and girls are matched on age, sex and pubertal stage were not confounded in our analyses; this aspect of our design may have aided our ability to detect sex differences in the association between INR and brain volume. Specifically, the effects of variation in INR may depend on ongoing neurodevelopment, which differs in boys in girls during puberty (Herting & Sowell, 2017). In addition to replicating the current findings, future studies should investigate the physical and hormonal changes associated with puberty that may underlie sexually dimorphic effects of INR on brain volume during adolescence.

We should point out five limitations of the current study. First, although this study was longitudinal, assessing adolescents at two time-points, we could not model the associations of INR with *trajectories* of brain volume, which may not be linear. Second, while a strength of TBM is the ability to simultaneously model effects across the whole brain, not just the cortex, TBM cannot distinguish effects from GM and WM when there are multiple tissue types in a given region, such as in areas of thin cortex. Thus, while we list the tissue type for each cluster in the tables, it is important to recognize that we list the *dominant* tissue type, not necessarily the only tissue type involved. Developmental processes differ in GM and WM, which can complicate interpretation; this inability to distinguish between tissue types, however, is attributable to current structural MRI standards and is not specific to TBM. Approaches that are purportedly tissue-specific (e.g., FreeSurfer [http://surfer.nmr.mgh.harvard.edu/] calculations of cortical thickness) may not necessarily be so. For example, for decades researchers have concluded that cortex thins over development, but recent evidence suggests that this is not necessarily the case; rather, increasing myelination alters tissue contrast, making it appear that cortex is thinning. Hopefully, future advances in MRI will allow investigators to characterize more accurately tissue-specific effects in GM versus WM. Third, our results may have been limited by our power to detect significant effects in certain analyses. Many of the effects we identified were “reversal” interactions (i.e., when associations are directly opposing within each level of a factor), and we do have greater power to detect such interactions; Giner-Sorolla, 2018). Additional studies with larger sample sizes are needed to determine whether these effects are replicable and whether smaller effects emerge when power is increased. Fourth, the measurements of INR in the current study were based on family income over the past 12 months and may not reflect children’s earlier environments. In this context, the timing of deficits or gains in family income may be an important moderator of the influence of family income on brain volume. As a related point, only 29% of the adolescents in the current study were low income relative to their community’s cost of living. Nevertheless, the fact that we observed associations of INR with brain volume even within this relatively higher SES sample suggests that normative variations in INR influence neurodevelopment. Finally, it is important to note that sex differences in the effects of INR on brain volume may be apparent at one developmental stage but not another(Joel & McCarthy, 2017), emphasizing the need to characterize the effects of variation in INR within specific developmental windows.

In closing, it is important to note that most research to date examining the effects of SES on neurodevelopment has spanned wide age ranges, has neglected to examine sex differences, and has used cross-sectional designs. Longitudinal investigations, such as the current study, facilitate the formulation of a comprehensive model for the effects of SES on neurodevelopment; however, the findings of the current study suggest that this model, perhaps not surprisingly, is quite complex. Future research is needed to replicate the findings of the current study and to investigate mechanisms for sexually dimorphic effects of INR on brain volume during adolescence. Although we replicated previous findings concerning the regions of the brain that are sensitive to variation in SES, we found that the direction of the associations between INR and volume differs depending on the region of the brain and on the basis of biological sex. Overall, biological sex may be an important variable to consider in analyses of the effects of SES on structural neurodevelopment across adolescence. In particular, boys appear especially sensitive to the effects of variation in INR on *change* in brain volume from early to later adolescence. Importantly, the current findings have implications for using neuroscience to inform policies to address socioeconomic disparities in child and adolescent health. Specifically, because markers of income-related risk may depend on biological sex, researchers should consider sex in designing strategies to identify vulnerable children and to prevent and mitigate the negative consequences of low family income.

## Disclosures and Acknowledgements

The authors report no conflicts of interest.

Funding for this study and support for authors was provided by the National Institutes of Health (R37 MH101495, U54 EB020403, R01 AG040060, R01 NS080655, K99 NS096116, and F32 MH107129); the Stanford University Precision Health and Integrated Diagnostics Center; the Brain and Behavior Research Foundation (Young Investigator Award 23819); the National Science Foundation Graduate Student Research Fellowship; Klingenstein Third Generation Foundation, and the Jacobs Foundation Early Career Research Fellowship (2017 1261 05). We thank Alexandria Price, Holly Pham, Isabella Lazzareschi, Cat Camacho, Monica Ellwood-Lowe, Sophie Schouboe, Maddie Pollak, Morgan Popolizio, Madelaine Graber, Anna Chichocki, Lucinda Sisk, Amar Ojha, Rachel Weisenburger, and Michelle Sanabria for their assistance in scheduling and running participants. We also thank the participants and their families for their contributions to this project.

## Footnotes

1 Family income-to-needs ratio takes into account the high cost of living in the San Francisco Bay Area communities from which we recruited our participants by adjusting family income by the low-income limit in the surrounding county corresponding to family size. Given wide variation in cost of living across the United States, the same income in different geographic areas may not reflect equivalent economic wellbeing. The average rent for an 815 sq. ft. apartment in the location of data collection is >$3,000/month (https://www.rentcafe.com/average-rent-market-trends/us/ca/santa-clara-county/palo-alto/).

2 We identified structures based on the coordinates of the cluster peaks; however, some voxels in the clusters labeled as *hippocampal cingulum* extended into hippocampal gray matter. TBM cannot distinguish effects on GM and WM in voxels that contain both.

