## Supplementary Text, Tables, and Analyses for "Cross-sectional and longitudinal associations of family income-to-needs ratio with cortical and subcortical brain volume in adolescent boys and girls" (from another version of this preprint: version 1, 2020-01-25)

**Measures**

**Severity of threat-related ELS**

Low socioeconomic status (SES) is associated with greater exposure to stressful life events (Evans & Kim, 2013). Therefore, it is possible that associations between lower family income-to-needs ratio (INR) and brain volume are explained by exposure to stressful life events. As previously described (King et al., 2017; King, Humphreys, Camacho, & Gotlib, 2019), at T1 participants were interviewed about their lifetime exposure to 30+ types of stressors using a modified version of the Traumatic Events Screening Inventory for Children (Ribbe, 1996). A panel of three coders, blind to the adolescents’ reactions and behaviors during the interview, then rated the objective severity of each type of stressor endorsed on a scale of (0 = non-event or no impact; 4 = extremely severe impact; ICC = 0.99). To quantify the severity of threat-related ELS, we summed the maximum objective severity ratings for family verbal and physical conflict, community verbal and physical conflict, abuse (physical, sexual, emotional), bullying, and exposure to war/terrorism (see <https://github.com/lucysking/els_stress_interview> for scoring script). We selected these events to be consistent with Sheridan and McLaughlin’s (Sheridan & McLaughlin, 2014) definition of threat as “the presence of an atypical (i.e., unexpected) experience characterized by actual or threatened death, injury, sexual violation, or other harm to one’s physical integrity” (p. 580). Scores for threat-related ELS ranged from 0.00-11.50. We found that INR and threat-related ELS were significantly negatively, but weakly, correlated (*r*(149)=-.26, *p*=.001), suggesting that these measures reflected different aspects of the environment in the current sample.

**Statistical Analyses**

**Main effects of INR on brain volume.** Although our primary analyses focused on examining sex differences in the association between INR and brain volume, we additionally tested the main effects of INR on brain volume. We conducted separate voxel-wise linear regressions testing the main effects of INR on brain volume, including sex, age, and race as covariates. The 9-DOF linear registration that is part of the TBM processing protocol accounts for differences in overall brain scale, removing much of the effect of intracranial volume (ICV); nevertheless, we also included ICV, computed from the linearly registered image, as a covariate. We present these models in equations 1 and 2:

Eq. 1	$X \sim A+ \beta1INR + \beta2Sex + \beta3Age + \beta4Race + \beta5ICV + \varepsilon$

Eq. 2 $\Delta X \sim A+ \beta1INR + \beta2Sex + \beta3Age + \beta4Race + \beta5ICV + \beta6Interval + \varepsilon$

where *X* is the jacobian determinant value at a given position, Δ*X* is the change in volume from early to later adolescence, *A* is the constant jacobian determinant term, the βs are the regression coefficients, and $\varepsilon$ is an error term. For each model, results were corrected for multiple comparisons across all voxels tested using Searchlight FDR (Langers, Jansen, & Backes, 2007). We report only those clusters that exceeded 50 voxels.

**Results**

**Cross-sectional and longitudinal main effects of INR on brain volume**

There were no significant main effects of INR on brain volume either cross-sectionally or longitudinally that survived correction for multiple comparisons. It is surprising that we identified no significant main effects of INR on brain volume. Although our sample size was large compared to some studies of the associations of SES with brain structure (e.g., Lawson et al., 2017; Mackey et al., 2015), several studies have implemented whole-brain analyses in considerably larger samples (e.g., Mcdermott et al., 2019; Noble et al., 2015). In these larger samples, main effects of SES on brain structure were small (β ~ 0.20). Our findings may have been limited by our power to detect such small effects. Importantly, for “reversal” interactions (i.e., when associations are directly opposing within each level of a factor), the power to detect an interaction is greater than the power to detect a main effect (Giner-Sorolla, 2018). It is possible that we would have detected main effects of INR on brain volume if we had conducted region-of-interest rather than whole-brain analyses. In previous analyses in this sample, however, we found no main effect of INR on hippocampal segmentations in early adolescence (King, Humphreys, Camacho, & Gotlib, 2019).

**Simple effects of INR on brain volume within each sex**

For each regression, we calculated studentized residuals to identify outlying observations using the “outlierTest” function in the R package “car” (Fox & Weisberg, 2019). For only three of the models did we identify observations with studentized residuals with Bonferonni-corrected *p*-values < .05. For the longitudinal model predicting change in volume of the right lingual gyrus, we identified one such observation (studentized residual=-3.85, Bonferonni *p*=.02). For the longitudinal model predicting change in volume of the right posterior thalamic radiation, we identified two such observations (first value: studentized residual=-4.02, Bonferonni *p*=.01; second value: studentized residual=-3.83, Bonferonni *p*=.02). Finally, for the longitudinal model predicting change in volume of the left superior frontal gyrus, we identified one such observation (studentized residual=6.14, Bonferonni *p*<.001). When we removed the outlying observations, results of the models were highly similar and conclusions were identical (i.e., effect sizes were of similar size and in the same direction). Therefore, the interaction effects do not appear to be driven by outlying observations.

Results of analyses in the age-matched subsample of boys and girls ages 11-12 years at T1 are presented in Table S1 and Figures S1 and S2.

**Table S1. Simple effect sizes of cross-sectional (A) and longitudinal (B) associations of INR in boys and girls in age-matched subsample**

| Structure (coordinates) | **Simple effect of INR**  **β (95% CI)** | |
| --- | --- | --- |
| A. Cross-sectional age-matched subset (N=53) | | |
| Positive in full sample | | |
|  | **Female** | **Male** |
| R. Cerebellar vermis (3, -47, -35) | -0.09 (-0.50, 0.33) | 0.60 (0.25, 0.95) |
| R. Midbrain (3, -32, -4) | -0.21 (-0.61, 0.19) | 0.26 (-0.09, 0.59) |
| R. Posterior thalamic radiation (40, -55, 2) | -0.16 (-0.56, 0.22) | 0.49 (0.16, 0.82) |
| R. Superior frontal gyrus (9, -8, 54) | -0.47 (-0.89, -0.05) | 0.37 (-0.01, 0.72) |
| R. Postcentral gyrus (15, -32, 69) | -0.06 (-0.48, 0.37) | 0.42 (0.06, 0.78) |
| R. Superior frontal gyrus (11, 46, 45) | 0.01 (-0.41, 0.44) | 0.52 (0.16, 0.88) |
| R. Middle frontal gyrus (51, 38, 5)^1^ | -0.12 (-0.56, 0.32) | 0.43 (0.06, 0.79) |
| L. Lateral occipital gyrus (-23, -89, 3) | -0.12 (-0.57, 0.32) | 0.35 (-0.03, 0.72) |
| L. Inferior frontal gyrus (-48, 46, -8) | -0.31 (-0.72, 0.11) | 0.48 (0.13, 0.83) |
| R. Superior temporal gyrus (63, -47, 19) | -0.08 (-0.48, 0.31) | 0.50 (0.17, 0.84) |
| Negative in full sample | | |
| L. Angular gyrus (-46, -39, 33) | -0.07 (-0.51, 0.35) | -0.12 (0.49, 0.24) |
| R. Thalamus (13, -23, 8) | 0.19 (-0.22, 0.61) | -0.28 (-0.64, 0.07) |
| L. Hippocampal cingulum/Fusiform gyrus (-20, -19, -33) | 0.31 (-0.11, 0.72) | -0.42 (-0.77, -0.07) |
| R. Hippocampal cingulum (23, -14, -23) | 0.55 (0.15, 0.95) | -0.26 (-0.59, 0.08) |
| L. Thalamus (-9, -24, 10) | 0.19 (-0.22, 0.61) | -0.28 (-0.64, 0.07) |
| L. Fusiform gyrus (-49, -49, -16) | 0.19 (-0.22, 0.61) | -0.31 (-0.68, 0.07) |
| B. Longitudinal age-matched subset (N=35) | | |
| Positive in full sample | | |
| L. Lingual gyrus (-20, -75, -5) | -0.28 (-0.81, 0.24) | 0.58 (0.24, 0.93) |
| L. Inferior temporal gyrus (-48, 1, -40) | -0.19 (-0.73, 0.35) | 0.77 (0.42, 1.13) |
| R. Lingual gyrus (12, -73, -9) | 0.52 (0.02, 1.00) | 0.65 (0.33, 0.98) |
| R. Superior parietal lobule (27, -60, 56) | -0.15 (-0.72, 0.42) | 0.64 (0.26, 1.02) |
| Negative in full sample | | |
| R. Posterior thalamic radiation (24, -70, 7) | 0.41 (-0.14, 0.96) | -0.71 (-1.08, -0.35) |
| L. Cerebellum (18, -66, -21) | 0.22 (-0.27, 0.71) | -0.84 (-1.16, -0.51) |
| R. Superior frontal gyrus (17, -5, 58) | 0.14 (-0.45, 0.73) | -0.50 (-0.89, -0.11) |
| L. Superior frontal gyrus (-16, 62, -10) | 0.13 (-0.34, 0.62) | -0.78 (-1.10, -0.46) |
| L. Superior longitudinal fasciculus (-33, -44, 16) | 0.27 (-0.27, 0.80) | -0.76 (-1.11, -0.40) |
| R. Hippocampal cingulum (26, -33, -7) | 0.37 (-0.18, 0.93) | -0.48 (-0.85, -0.11) |

**Notes. “**Positive/negative in full sample” indicates whether the interaction was positive/negative in the full (non-age-matched) sample. “Positive” interactions indicated that the association between INR and volume was significantly more positive in boy than in girls; “negative” interactions indicated the association between INR and volume was significantly more negative in boys than in girls.

**Figure S1. Scatterplots of results from cross-sectional INR ×** **sex analysis in the subsample of 53 age-matched boys and girls**

A. Positive in full sample

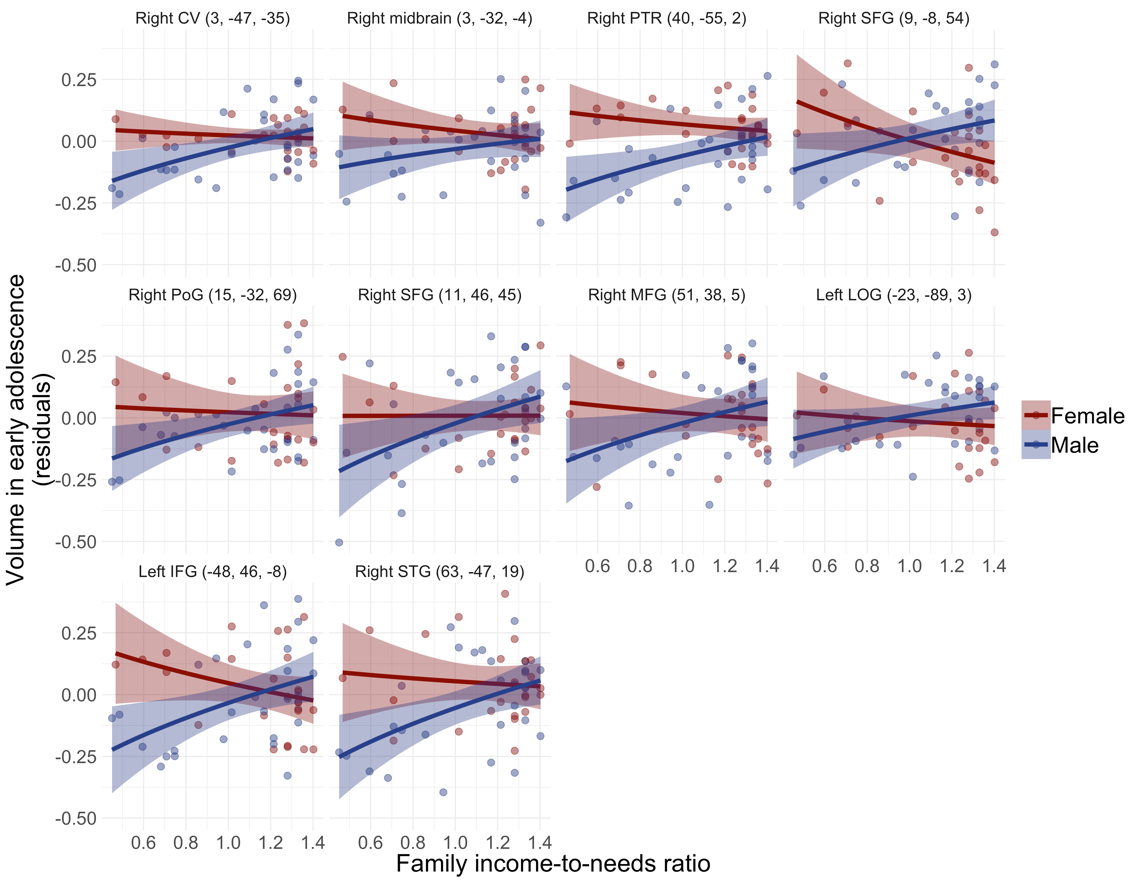

B. Negative in full sample

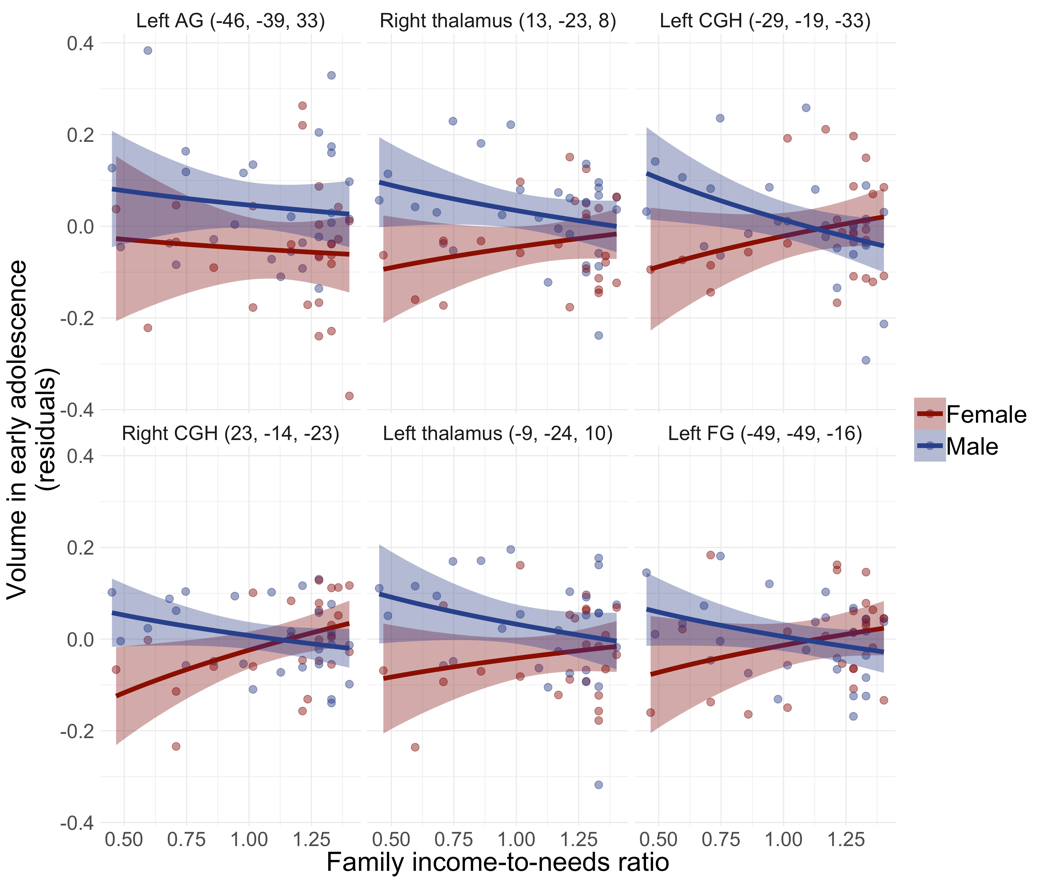

**Notes.** Coordinates of cluster peaks are in parentheses. Positive clusters (A): CV = cerebellar vermis, WM = white matter, PTR = posterior thalamic radiation, SFG = superior frontal gyrus, PoG = postcentral gyrus, MFG = middle frontal gyrus, LOG = lateral occipital gyrus, IFG = inferior frontal gyrus, STG = superior temporal gyrus. Negative clusters (B): AG = angular gyrus, CGH = hippocampal cingulum, FG = fusiform gyrus.

**Figure S1. Scatter plots of results from longitudinal INR ×** **sex analysis in the subsample of 35 age-matched boys and girls**

A. Positive in full sample

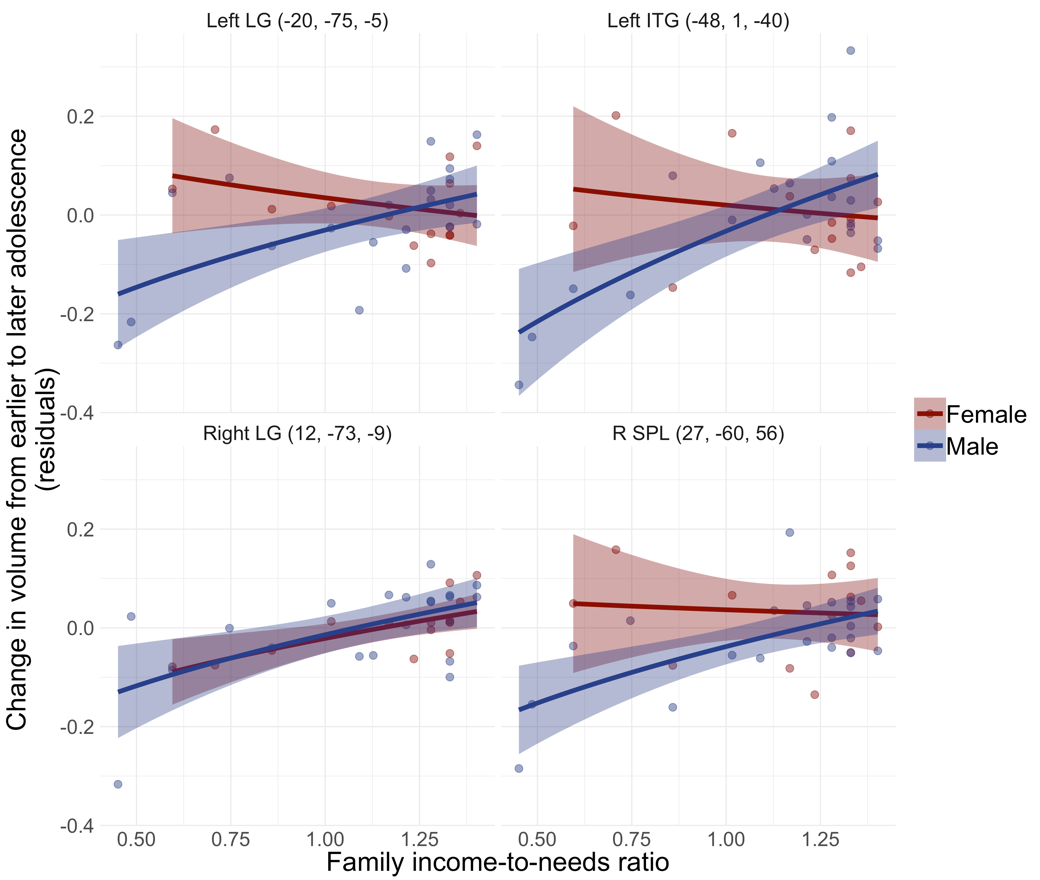

B. Negative full sample

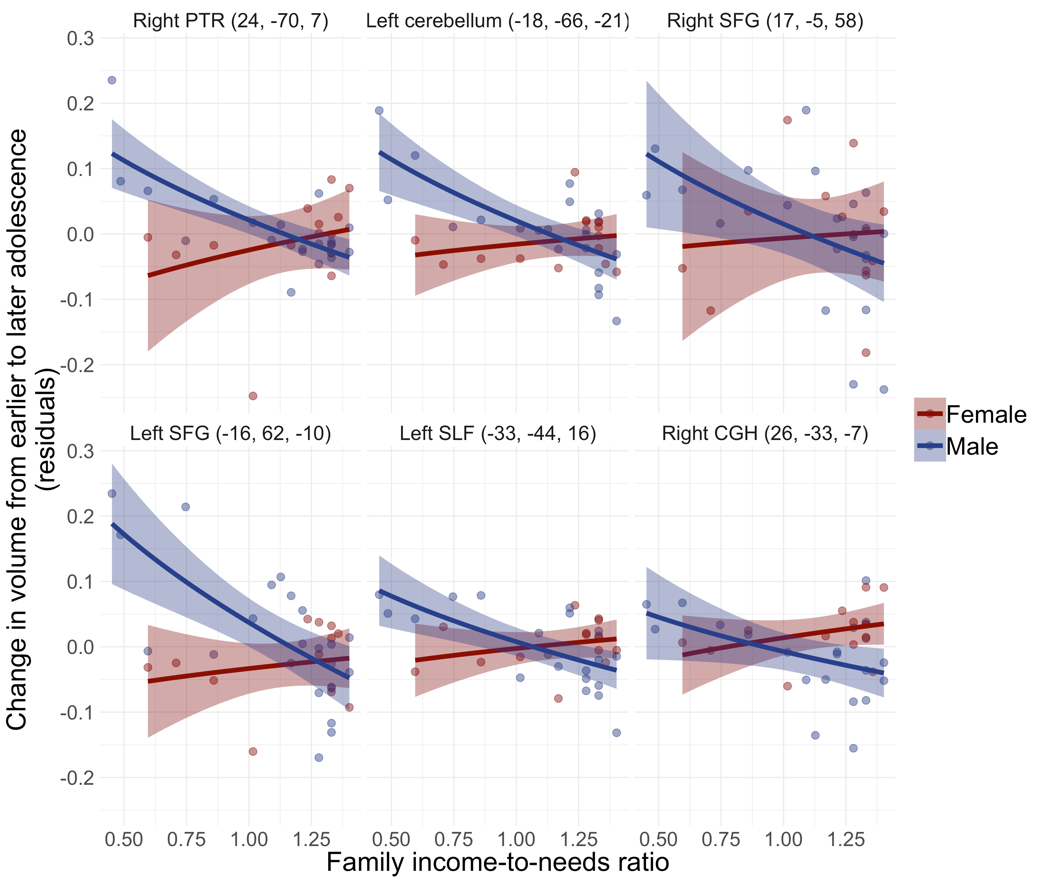

**Notes.** Coordinates of cluster peaks are in parentheses. Positive clusters (A): CV = cerebellar vermis, WM = white matter, PTR = posterior thalamic radiation, SFG = superior frontal gyrus, PoG = postcentral gyrus, MFG = middle frontal gyrus, LOG = lateral occipital gyrus, IFG = inferior frontal gyrus, STG = superior temporal gyrus. Negative clusters (B): AG = angular gyrus, CGH = hippocampal cingulum, FG = fusiform gyrus.
